# A hypothesis-driven approach to assessing significance of differences in RNA expression levels among specific groups of genes

**DOI:** 10.1101/136143

**Authors:** Mingze He, Peng Liu, Carolyn J. Lawrence-Dill

**Author notes:** communications (ORCID 0000-0003-0069-1430).

## Abstract

Genome-wide molecular gene expression studies generally compare expression values for each gene across multiple conditions followed by cluster and gene set enrichment analysis to determine whether differentially expressed genes are enriched in specific biochemical pathways, cellular components, biological processes, and/or molecular functions, etc. This approach to analyzing differences in gene expression enables discovery of gene function, but is not useful to determine whether pre-defined *groups* of genes share or diverge in their expression patterns in response to treatments nor to assess the correctness of pre-defined gene set groupings. Here we present a simple method that changes the dimension of comparison by treating genes as variable traits to directly assess significance of differences in expression levels among pre-defined gene groups. Because expression distributions are typically skewed (thus unfit for direct assessment using Gaussian statistical methods) our method involves transforming expression data to approximate a normal distribution followed by dividing the genes into groups, then applying Gaussian parametric methods to assess significance of observed differences. This method enables the assessment of differences in gene expression distributions within and across samples, enabling hypothesis-based comparison among groups of genes. We demonstrate this method by assessing the significance of specific gene groups’ differential response to heat stress conditions in maize.

**Abbreviations:**

GO: – gene ontology HSP – heat shock protein
KEGG: – Kyoto Encyclopedia of Genes and Genomes
HSF TF: – heat shock factor transcription factor
HSBP: – heat shock binding protein
RNA: – ribonucleic acid
TE: – transposable element
TF: – transcription factor
TPM: – transcripts per kilobase millions

## 1. Introduction

Determining gene function remains a fundamental problem in biology. Measuring gene expression levels via transcript analysis across various treatments and developmental stages from many tissues greatly facilitates gene, pathway, and genomic functional annotation and interpretation. Because gene expression measurements are highly variable, many existing sophisticated statistical models and implementations are available to reduce measurement bias introduced during sampling and technical procedures^1,2,3,4,5,6,7^. After these calibration and normalization procedures, researchers commonly work toward identifying a set of differentially expressed genes then analyze annotations to those genes in an effort to discover shared functions (based on, e.g., shared biochemical pathways or biological processes; see Figure 1). Existing methods^8,9,10^ for such analyses are highly dependent on the accuracy and precision of gene function annotations from gene ontology (GO) terms^11^, KEGG pathway enzyme annotation^12^, etc.^13^ In other words, enrichment analysis depends on reliable functional annotation. Because some annotations are based on, e.g., experimental evidence whereas others derive from computational prediction alone,^13^ annotation bias, incorrect annotations, and low quality annotations could lead to misinterpretation of detected expression patterns.

**Figure 1.**
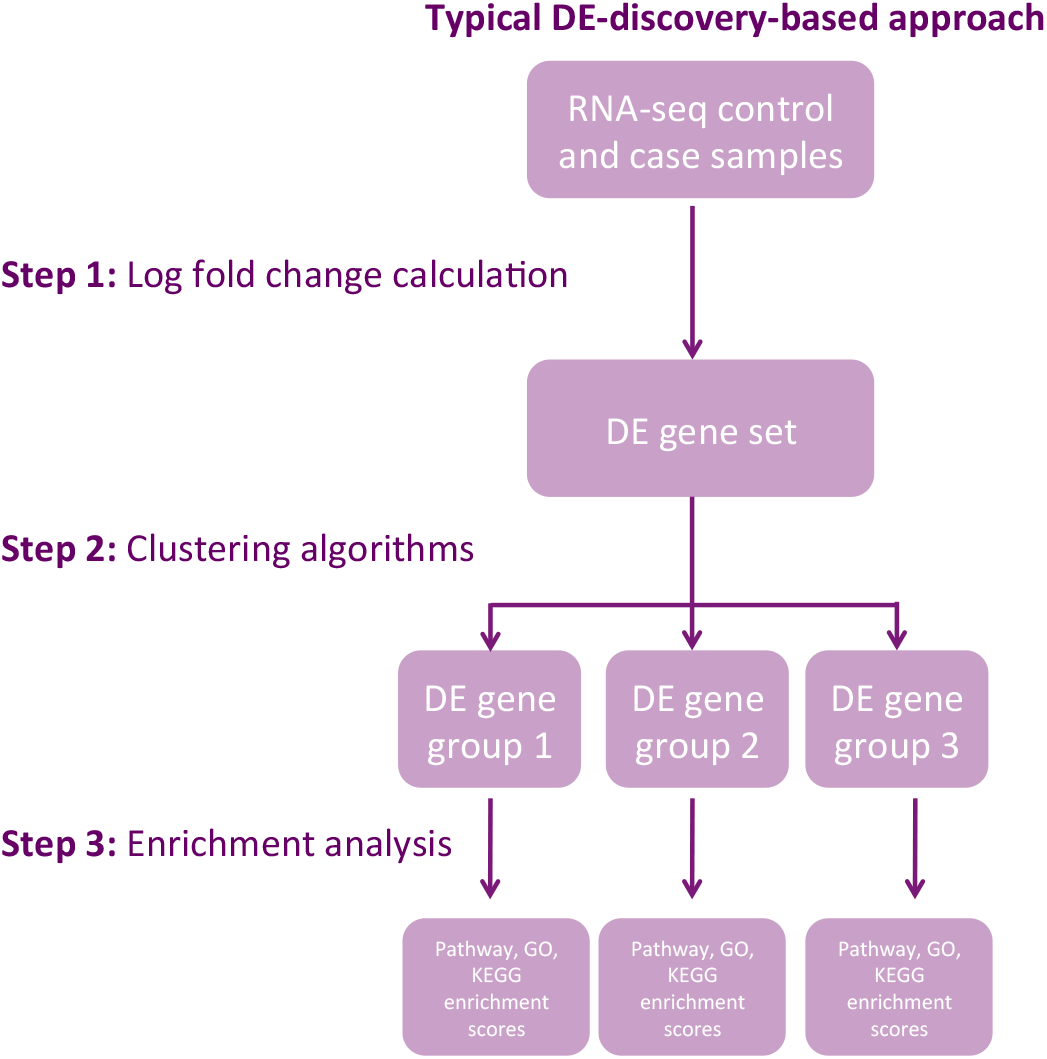
Discovery-based approach to gene expression analysis. Boxes represent data or analysis results, arrows represent analyses. Step 1: RNA-seq samples are subjected to log fold change analysis to identify a DE gene set. Step 2: clustering algorithms are used to create DE clusters (groups 1, 2, and 3). Note that step 2 maybe skipped, especially when the list of DE genes is not long. Step 3: Enrichment analysis identifies pathways, GO terms, KEGG terms, etc. associated with the DE gene groups.

Other, less commonly used approaches assess differential expression within predefined gene sets^14,15^. Such approaches are either ‘competitive’ or ‘self-contained,’ terms coined by Goeman and Buhlmann^2^. The competitive approach identifies a gene set with comparatively greater or comparatively fewer differentially expressed genes relative to all other genes in the dataset. The null hypothesis for the competitive approach is as follows: “Differential expression in a defined subset of genes is observed, at most, as often as it is observed for genes in the remaining set.” The self-contained approach focuses only on the information from gene sets of interest. The null hypothesis underlying self-contained tests could be stated as, “No genes in the defined subset are differentially expressed.” Each approach has important caveats. Competitive group analysis depends on the background distribution and assumes independent sampling^2,16,17^. Self-contained analyses can be highly affected by a very small number of highly expressed genes; thus one highly expressed gene could result in failure to detect otherwise significant patterns. Another drawback to these methods is the fact that enrichment analyses, designed to test whether specific gene groups are either up- or down-regulated under specific conditions enriched in certain biological functions, do not produce a direct measurement of how *much* gene group expression levels change across experiments.

Here we present and describe a different method to comparatively assess the expression patterns among pre-defined groups, both within and across treatments. The key difference between existing methods (Figure 1) and ours (Figure 2) is the absence of a ‘differentially expressed gene set’ concept throughout the process. We treat each normalized gene expression value as a variable in the later comparison between groups. Our method enables examination on non-linear, non-one-to-one correspondence among gene groups.

**Figure 2.**
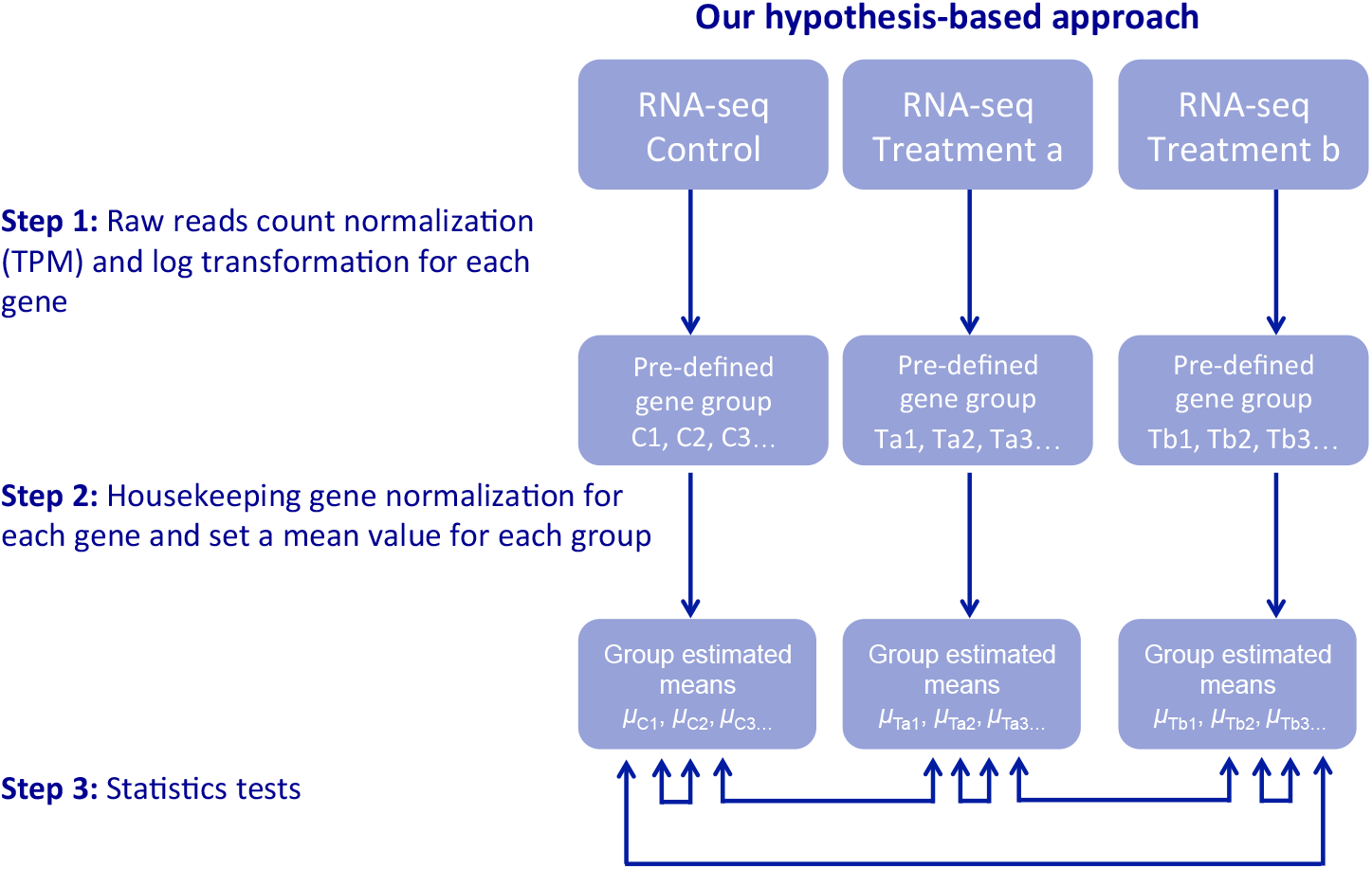
Our hypothesis-based approach to gene expression analysis. Boxes represent data or analysis results, arrows represent analyses. Step 1: RNA-seq samples (control and treatment) are subjected to raw read count normalization (transcripts per kilobase million; TPM) and log transformation for each gene. Step 2: Pre-defined gene groups (Control 1, Control 2, Control 3, Treatment a1, Treatment a2, etc. where the gene list for C1 is the same as Ta1 and Tb1, etc.) are defined and housekeeping genes are designated for normalization and to create group estimated means. Step 3: Gaussian statistical tests are carried out within (e.g., *μ*_Ta1_ and *μ*_Ta2_) and/or across samples (e.g., *μ*_Tb2_ and *μ*_Tb2_).

We demonstrate this new method by assessing a well-studied biological phenomenon: the response of genes to heat stress. Heat shock proteins (HSP) are regulated by heat shock factors (which are a specific group of transcription factors; abbreviated here HSF TFs) under heat stress^18,19^. HSF TFs are negatively regulated by heat shock factor protein bindings (HSPBs)^20^. By dividing maize genes into subgroups, i.e., HSPs, HSPBs, HSFs, other TFs, and housekeeping genes, we compare each group’s response to heat stress.

We also show the method’s utility for analyzing groups of genes that are not functionally related by evaluating the significance of differences in gene expression for maize genes nearby *gyma* and *huck* transposable elements (TE) in heat stress conditions, confirming Makarevitch et al.’s findings that genes with *gyma* nearby are significantly up-regulated whereas genes with *huck* nearby show no differential expression in heat stress conditions^21^.

## 2. Materials and Methods

For the first example, we focused on these subgroups of maize genes: HSPs (reported in Pegoraro *et al*.^22^), HSPBs (from Gramene^23,24^), HSF TFs and other TFs (from GRASSIUS^25,26^), and housekeeping genes (see “Supplementary Material 2” from Lin *et al*.^11^). Each group’s response to heat stress was compared across conditions and across subgroups within the same condition using RNA-seq datasets reported by Makarevitch *et al*.^21^ Gene groups derived from these sources are listed in Supplementary Table S1 (all Supplementary Materials are available online at DOI 10.17605/OSF.IO/Y46TF, which currently resolves to https://osf.io/y46tf/).

For the second example, we use the Makarevitch *et al*.^21^ expression dataset along with their TE gene set lists and coordinates from the maize TE dataset reported by Baucom *et al*.^27^ where TEs are considered ‘nearby’ if they are +/-1 kb from the transcription start sites of genes in the B73 RefGen_v3 genome assembly with gene annotation version ZmB73_5b^28^. We focus on genes near *gyma* TEs, which constituted the largest set of genes near a specific TE type that were up-regulated by heat stress, and compare those to genes near *huck*, which constituted the largest subset of genes near a TE family that were not differentially expressed. Gene groups derived from these sources are listed in Supplementary Table S2.

Because this method aims to detect the differences between groups of genes based on a hypothesis-driven approach, methods for grouping genes are not limited to genes with homologous sequences (such as groups in the first example) or shared pathways, though gene groups assembled using such concepts are reasonable candidates for this approach. One example of a non-homologous gene set made up of members that also are not necessarily involved in shared biochemical pathways includes members of the housekeeping gene set (such as those used in the normalization step described in Theory/Calculations), which we define as those genes essential to maintaining basic cellular functions and including only those with constant expression values across various tissues and conditions. Other groups of genes share such characteristics as having a TE located near the transcription start site (the second example), genes in a particular region of a chromosome, or genes sharing structural elements. These groups each share some characteristic, but these characteristics are not limited.

## 3. Theory/Calculations

Suppose we have a pre-defined gene set, for which read counts for each gene have been normalized by TPM that is calculated as a percentage of reads normalized by length of each gene to total reads within a sample^29^. Because we intend to treat each gene as a variable, we use the TPM value averaged across biological replicates to measure the expression level for each gene:

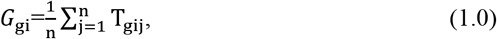

where T_gij_ stands for the TPM value of the g^th^ gene for the j^th^ biological replicates within the i^th^ condition/treatment, j = 1,2,…, n_i_, and n_i_ is the number of replicates for the i^th^ condition. This step summarizes expression levels for each gene to facilitate the comparison using gene sets.

We apply the logarithm transformation as in (1.1) because the expression measurements are typically highly skewed to the right. We add a small value 1 to each TPM to avoid problems for the logarithm transformation.

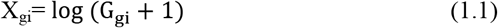

Secondly, we calculate a normalization factor based on housekeeping genes:

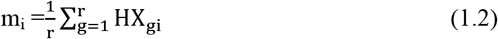

where HX_gj_ are expression values of the g^th^ housekeeping gene in the i^th^ condition after transformation using (1.0) and (1.1), and suppose that there are a total of r housekeeping genes.

Because housekeeping genes are, by definition, expected to express at a constant level across tissues and under various conditions, we could use mi to normalize across conditions (X_gi/mi_) when we compare a gene set from different samples.

Finally, Student’s *t*-test (or other appropriate parametric or non-parametric tests) could be applied to test two different types of null hypotheses.

First, we can test for equal means between gene groups A and B under the same treatment condition, e.g., condition 1 (1.3).

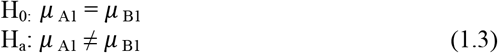

This might be of interest to determine, e.g., whether a transcription factor and the genes it activates respond similarly in a given condition.

Second, we could also test whether the same gene set (A) has the same mean across conditions 1 and 2:

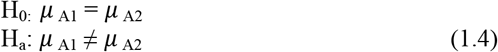

This might be of interest to determine, e.g., whether a group of genes with increased transcript abundance under salt stress conditions shows similar increases in transcript abundance in drought stress conditions.

In both examples, if a gene has an average TPM value (*G_gi_*) below 1 in both conditions, we consider such data unreliable because it is hard to tell whether the corresponding reads were from signal or noise. Therefore we exclude such genes from the following analysis. Note that we used TPM in the above formulae, but FPKM values can also be used.

## 4. Results and Discussion

As is often observed for gene expression values, the distribution of expression values across all genes in shoot tissues from maize seedlings follows an exponential curve, with the majority of genes expressed at relatively low levels (Figure 3a; non-stressed condition). Log transformation results in a distribution much closer to normal (Figure 3b). Log-transformed data collected from the shoot tissues of maize seedlings under both non-stress (Figure 3c) and heat stress conditions (Figure 3d) generally follow the normal distribution (i.e., 93.1%-97.4% of all log transformed data were located within a 95% confidence interval), indicating that this transformation approach is reasonable. Note well: because our method relies upon transformation to approximate a normal distribution, it is important to check the results of log transformation not only for all sampled genes and for all conditions, but also for each individual gene group and treatment combination. In this example, the housekeeping genes, HSFs, other TFs, and HSPBs all appear to roughly approximate the normal distribution (as assessed via QQ-plot; Figs. S1-8). However, the transformed expression patterns for HSPs do not follow the normal distribution (see Figs. S9 and S10) and are therefore not appropriate for analyses using parametric tests of significance among groups. This result exemplifies the need to inspect transformed distributions as a step in applying this method.

**Figure 3.**
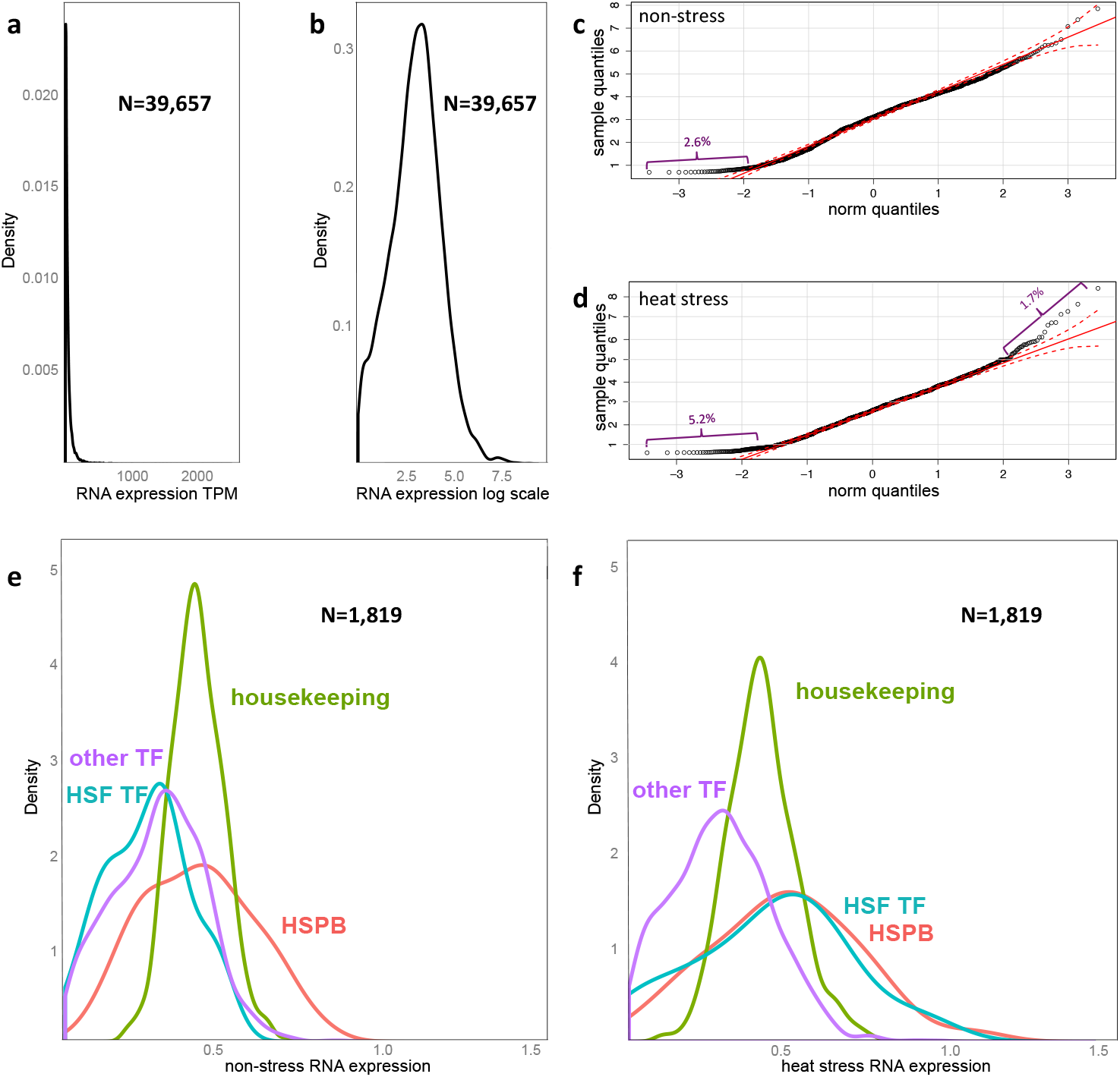
Log transformation enables Gaussian modeling of expression patterns among groups of genes. (**a**) The density plot of all maize genes (N=39,657) with a given RNA expression level is plotted for the non-stress condition. (**b**) Log transformation of RNA expression values before m normalization results in a roughly normal distribution. (Note y axis is not the same between panels a and b.) A QQ-plot (normal distribution quantiles plotted against sample quantiles) for the log-transformed data collected from (**c**) non-stress and (**d**) heat stress shown as black circles. Red diagonal indicates perfect concordance. Red dashed lines indicate the 95% confidence interval (CI). Percentage of data falling outside the 95% CI are indicated in purple. (**e** and **f**) RNA expression levels of HSF TF, HSPB and other TF gene groups normalized by housekeeping genes are plotted by percentage. Housekeeping genes shown in green, HSPB in red, HSF TFs in turquoise, and other TF family genes in purple. (**e**) Non-stress condition. (**f**) Heat stress.

As one might expect given the known and well-understood biology of response to heat stress, transcription of HSF TFs increases in response to heat stress and shows a very different distribution than other TFs (Figure 3 panels e and f). Relative to the non-stress condition, HSF TF s have right-shifted RNA expression distributions relative to housekeeping genes and other TFs under heat stress. Beyond inspecting the distributions, this data transformation approach allows application of parametric statistical approaches, e.g., the t-test, to compare mean values between distributions within a given sample. As shown in Table 1, under non-stress conditions, the t-test fails to reject the null hypothesis (i.e., HSF TF and other TF have no differences in mean values). However, as shown in Table 2, under heat stress t-test results reject the null hypothesis, indicating that the higher expression of HSF TFs is significantly different than that of other TFs. As shown in Table 3, the expression distribution shifts between Figure 1 panels e and f are significant only for HSF TFs, but not for other TFs nor for the HSPB group.

**Table 1.** *t*-test p-values between gene sets under non-stress conditions.

|  | Sample size | HSF TF | HSPB | Other TF |
| --- | --- | --- | --- | --- |
| HSF TF | 19 | - | <0.0004* | 0.272 |
| HSPB | 37 | - | - | 0.0001* |
| Other TF | 1,299 | - | - | - |

\* p-values smaller than 0.05 after Bonferroni multiple test correction.

**Table 2.** *t*-test p-values between gene sets under stress conditions.

|  | Sample size | HSF TF | HSPB | Other TF |
| --- | --- | --- | --- | --- |
| HSF TF | 23 | - | 0.5003 | 0.0060* |
| HSPB | 37 | - | - | <0.0001* |
| Other TF | 1,234 | - | - | - |

\* p-values smaller than 0.05 after Bonferroni multiple test correction.

**Table 3.** *t*-test p-values among the same groups of genes under two conditions.

|  | HSF TF | HSPB | Other TF |
| --- | --- | --- | --- |
| Heat vs normal | 0.0039* | 0.1659 | 0.2457 |

\* p-values smaller than 0.05 after Bonferroni multiple test correction.

It is worthwhile to show that the approach we have developed is versatile beyond the analysis of homologous genes with shared functions given that hypotheses concerning concerted action of genes can be made for various groups (e.g., genes that share specific TF binding sites, genes that all participate in the same biological pathway, etc.). For our second example, we investigated differences in expression patterns among genes near TE types that Makarevitch *et al*. showed may affect transcription under heat, cold, high UV, and salt stress conditions^21^. We analyzed their maize RNA-seq dataset collected under normal and heat stress using our new method as described in the first example. Among other findings, Makarevitch *et al*. showed that 25% of genes with a *gyma* element +/-1 kb of the transcription start site are up-regulated under heat stress whereas those with the *huck* element in the same location are not^21^. We plot transformed expression patterns of groups of genes with the *gyma* and *huck* motifs near the transcription start site (as well as housekeeping genes) under nonstress and heat stress scenarios (Figure 4). Those plots show that the distribution of genes with *gyma* near the transcription start site shifts to the right under heat stress conditions, which indicates a positive transcriptional response to heat stress. Genes with *huck* near the transcription start site do not. We then apply our method to assign p-values to assess the significance of both *gyma* and *huck* to the activation of nearby gene expression (for QQ-plots see Figs. S11-S14). The *gyma* and *huck* distribution means are significantly different from each other in non-stress conditions (Table 4), but in heat stress conditions the means of *gyma* and *huck* expression distributions are not significantly different from each other (Table 5). Compared across samples (non-stress *vs* heat stress), the expression differences for the *gyma* gene group means yields a p-value < 0.0001 (see Table 6). This indicates a statistically significant positive response to heat stress for genes with *gyma* nearby. Genes with the *huck* TE family nearby show no significant changes in the distribution of transformed expression values (p-value=0.76). These results not only confirm the findings of Makarevitch et al., they assign significance to the observation that genes with *gyma*, but not *huck*, near the transcription start site show increased transcription under heat stress^21^.

**Figure 4.**
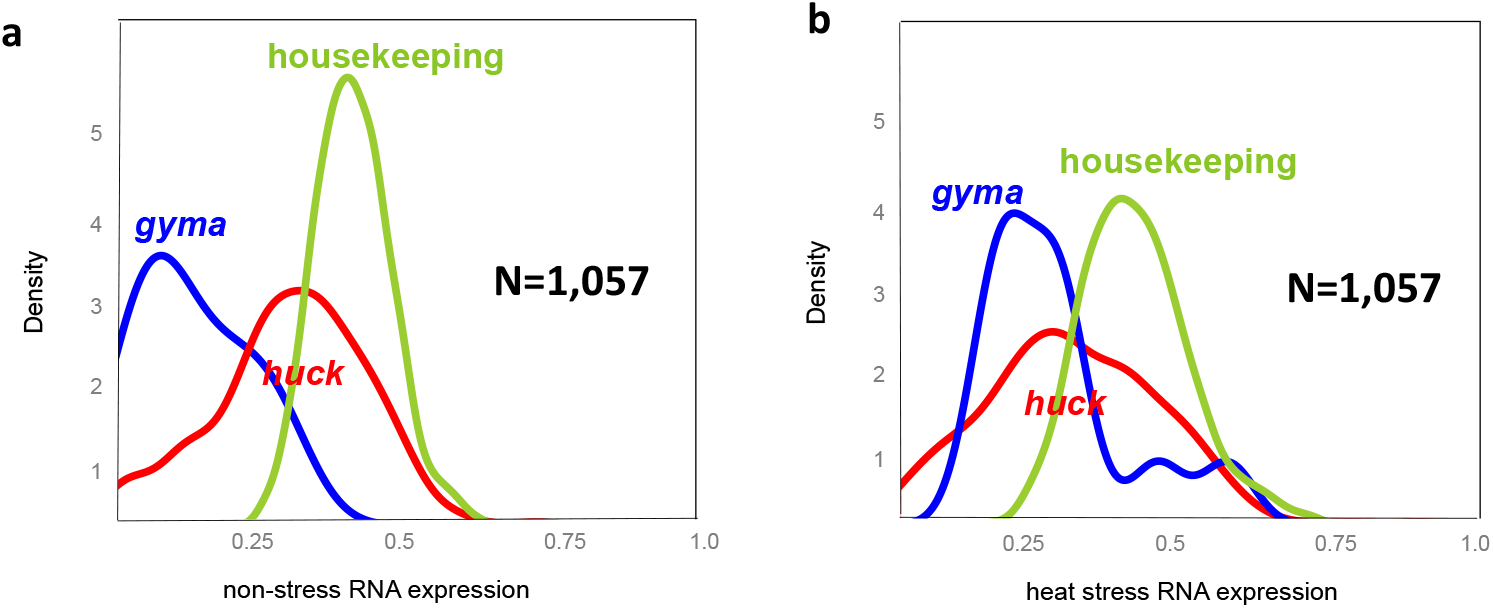
TE within and near the promoter region activate nearby gene transcriptional response to heat stress. RNA expression levels of gene groups with *gyma* or *huck* near the promoter region normalized by housekeeping genes are plotted by percentage. Housekeeping genes shown in green, *gyma* in blue, and *huck* in red. (**a**) Non-stress condition. (**b**) Heat stress.

**Table 4.** *t*-test p-values between gene sets under non-stress conditions.

|  | Sample size | *gyma* | *huck* |
| --- | --- | --- | --- |
| *gyma* | 11 | - | <0.0001* |
| *huck* | 234 | - | - |

\* p-values smaller than 0.05 after Bonferroni multiple test correction.

**Table 5.** *t*-test p-values between gene sets under stress conditions.

|  | Sample size | *gyma* | *huck* |
| --- | --- | --- | --- |
| *gyma* | 11 | - | 0.4640 |
| *huck* | 213 | - | - |

**Table 6.** *t*-test p-values among the same groups of genes under two conditions.

|  | *gyma* | *huck* |
| --- | --- | --- |
| Heat stress vs non-stress | <0.0001* | 0.7575 |

\* p-values smaller than 0.05 after Bonferroni multiple test correction.

## 5. Conclusions

We changed the dimension of comparison for gene expression studies to derive a new hypothesis-driven approach to gene expression analysis that enables the use of Gaussian parametric tests to assess results. One could also easily use this approach to study other phenomena and to test various biological hypotheses. For example, recent studies reporting gene regulation show that motifs around regulatory regions of genes (e.g., transposable elements^21^, high GC content motifs^30^, and G-quadruplexes^31^) may influence nearby gene expression levels under stress conditions. One could also apply this approach to evaluate the influence of gene sequence composition bias on expression (e.g., GC content effects) as well as expression differences that may be attributable to a gene’s local context (e.g., location on the chromosome or adjacency to other genes).

By design, our approach is limited to pre-defined gene groups. Thus, this method is not suitable to discover genes that respond to, e.g., a particular stress environment, which is often the goal of a typical discovery-based approach to gene expression analysis. Also, our approach does not attempt to assess the correctness of gene groupings. For example, if some HSF TF had been missing from the HSF TF gene group, the analysis could still yield significant p-values. On the other hand, the power of this method is that it can be combined with existing knowledge about specific groups of genes to assess how well a given annotation method grouped genes based on shared gene expression patterns.

This new method is complementary to existing expression analysis tools such as those available via the Gramene Plant Reactome^32^. First, our method is not limited to homologous genes or pathway membership, thus, we broaden the scope of comparison that can be made. Second, our distribution-based visualization is a graph-based alternative to heatmap or color-coded plots used in pathway analysis, which can be difficult to interpret (especially for individuals with some types of colorblindness). Third, our method allows for statistical tests among groups of genes rather than between individual genes, where individual gene comparisons may conflict and cause confusing results for downstream analyses including, e.g., pathway analysis. Our test could be used to eliminate such a conflict by transforming each gene considered into a variable, and all genes in a certain pathway into a group.

Novel approaches to enable hypothesis testing for shared regulation of gene sets are needed. As technologies for measuring gene expression improve, we anticipate that additional methods will be developed that expand the toolset for grouping-oriented gene expression analytics.

## Acknowledgements

Funding in support of this work came from the Iowa State University Plant Sciences Institute (CJLD and MH) and NSF PGRP #1238142. We thank Irina Makarevitch at Hamline University for providing TE annotation files and for review. We thank Jennifer Clarke at University of Nebraska – Lincoln, Drena Dobbs and Justin Walley at Iowa State University, and members of the Lawrence-Dill Plant Informatics and Computation Lab for suggestions and helpful discussions.

